# Cyanobacteria is uniquely enriched in the roots of grain amaranths

**DOI:** 10.1101/540484

**Authors:** Suran Nambisan, Meeta Sunil, Bibha Choudhary, Subhashini Srinivasan

## Abstract

Plants and microbes have coexisted for millions of years and some have evolved mechanisms to achieve symbiosis driving positive selection. The bartering of chemicals produced by soil microbes and plants favour enrichment of certain types of bacteria near the roots that offers selective advantage to the plant under a given environment. Grain amaranths display certain important agronomic characteristics like C_4_ dicot, high protein and high lysine grains, resistance to biotic and abiotic stress, which can be translated to other crops. Considering an unusual collection of desirable traits shown by grain amaranths, it is worth pondering if symbiosis with bacteria has played any role in these traits. The objective of this study is to identify bacterial root microflora unique to grain amaranths. Here, by comparing rhizospheric and endophytic composition of 16S rRNA from various sections of roots from selected species under major plant orders including the three varieties of grain amaranths, we report that Cyanobacteria are uniquely enriched by grain amaranths. The diversity in OTUs among the Cyanobacteria also significantly increased among samples from amaranth species compared to negative control. This finding is also validated using root transcriptome of *Amaranthus hypochondriacus*, where we observe relative increase in Cyanobacterial population between day 15 to day 30 compared to other abundant phylum during this period.

## INTRODUCTION

Plants have evolved to survive in diverse ecological niches with help of specific adaptations, some involving symbiotic associations with a few of the myriad microorganisms present in soil. The nature and composition of soil [1], the microbes residing in it [2] and the genotype of the host plant [3,4] contribute to the selection of helpful microbes seen in such interactions which are required for successful adaptation by plants in environments like hypersalinic conditions [5]. A closer look at the rhizospheric and endophytic microbes associated with the plant roots can reveal interesting facets about plant-microbial interactions, which play critical roles in the survival and desirable traits of plants. One of the most well documented plant-microbe symbiosis is between legumes like *Medicago truncatula* [6] and soybean [7] and rhizobia, which are involved in nitrogen fixation [8]. A few important crop species from the family Poaceae like rice (*Oryza sativa*) have also been keenly studied for their symbiotic associations with endophytic microbes like rhizobia [9]. Another study in finger millet (Poaceae family) has shown a root-inhabiting bacterial endophyte to help fight an invading fungal pathogen [10]. Several other studies involving plant-microbe symbiotic associations describe the ability of plant species to develop abiotic stress tolerance [11] like salt [12], drought [12] and as well as pathogen resistance [13, 14, 15] with the help of associated microbes. For example, inoculation of *Pseudomonas* sps. on wheat seedlings increase root growth [16] and bacterium *Stenotrophomonas rhizophila* strongly increases drought tolerance in crops like sugar beets and maize [11].

Until a few years back, an in-depth study of the microbes involved in plant-microbe symbioses had been hampered by the limited range of methods available for culturing bacteria that can be characterized and studied. The advent of high-throughput sequencing technologies has now allowed us to study plant-microbe symbiosis at an unprecedented scale and depth.

Metagenomics and metatranscriptomics are increasingly being used to survey entire microbial communities like the human gut [17], soil [18], sludge [19] and water bodies [20]. In recent times, several groups have employed these high-throughput techniques to understand the microbial community structure associated with plant roots and how variations in the root microbiota and host genotype affect each other. Many plant species belonging to taxonomically diverse groups have been surveyed for their microbial communities, ranging from model organisms like *Arabidopsis thaliana* [21, 22, 23, 24, 25] (fam. *Brassicaceae*), *Lotus japonicas* [26] and *Medicago trunculata* [27] (fam. *Leguminosae*) to important crop species, like soybean [28] (fam. *Leguminosae*), monocots like rice [29,30], wheat [28], maize [31,32] and barley [33] (fam. *Poaceae*), sugarbeet [34,35] (fam. *Amaranthaceae*), tomato [36], potato [37] (fam. *Solanaceae*) and wild plants and trees like *Boechera stricta* [38] (fam. *Brassicaceae*) and *Populus deltoides* [39] (fam. *Salicaceae*).

Grain amaranth comes under the plant order Caryophyllales, which is sparse in edible plants and has not been widely studied using genomics techniques. More recently, the draft genome and transcriptome of two varieties of *Amaranthus hypochondriacus* has been deciphered [40, 41, 42]. This resource is critical for metagenomic and metatranscriptomic interrogation of amaranth roots. Here we have interrogated the roots of three grain amaranths species for the enrichment of unique microbial species.

## RESULTS

For this study, metagenomic DNA was isolated from three types of samples i.e., root (R), rhizospheric soil (RS) and surrounding outer rhizospheric soil (S) for three varieties of grain amaranths (*A. hypochondriacus, A. cruentus* and *A. caudatus*), and three species under other major plant orders viz., *Beta vulgaris (Caryphyllales), Cicer arietinum* (Rosid) and *Solanum lycopersicum* (Asterid). Bulk soil was collected as soil sample (SOIL) for control from the same plot. Only two fractions (R and S) were taken for wild amaranth plants used in the study. Metagenomic DNA was isolated for these samples were also isolated as well. In all, this amounts to the 21 samples sequenced in the study.

Isolated DNA was subjected to 16S rRNA amplicon sequencing of the V3 region using Illumina MiSeq platform for profiling the bacterial microbiota. Each species was sequenced to a depth of 0.5 million paired-end reads per sample and a read length of 150 bases. The data was filtered and analyzed using QIIME pipeline as described in the method section.

### Sequencing Analysis

One hundred and fifty million bases per sample of 16S rRNA V3 amplicon sequencing was done using Illumina MiSeq platform for the 21 samples. Overall 13,275,951 raw reads were generated of which more than 85% were with a Phred quality score above Q30. The raw reads were filtered using FastQC [43] and trimmed before assembling reads into full-length amplicons using PANDAseq [44]. We got a total of 11,207,468 amplicons with an average depth of 533,689 reads per sample and mean length of 148.69 +/-11.61 bases. A summary of the reads for samples is given S1 Table.

### OTU Analysis

In all, 591,444 OTUs were identified using the QIIME pipeline [45] for all samples. All the identified amplicons belonged to Eubacteria and no OTUs were derived from Archaea or Eukarya. After removal of chimeric OTUs and singletons the number of OTUs for all samples was 134,362 (observations/sample: min=2396, max=47612, mean=20824.6, std. dev.=15468.2) and total counts for all OTUs was 9890033 (counts/sample: min=271356, max=556774, mean=470953.9, std. dev.=71685.7). The top 10 phyla in terms of percent OTUs and abundance based on number of amplicons are represented in the Fig 1 and Fig 2 respectively. Overall the dominant phyla with the largest number of OTUs across all samples were Actinobacteria and Proteobacteria followed by Firmicutes, Chloroflexi, Acidobacteria and Bacteroidetes (Fig 1). In amaranth samples OTUs from Actinobacteria were predominant than Proteobacteria. However, as shown in Fig 2, most abundant OTUs based on percentages of amplicons representing the OTUs, show very high abundance of Cyanobacteria uniquely in samples from amaranth species. For example, although the diversity in OTUs from Cyanobacteria is relatively smaller compared to Proteobacteria and Actinobacteria (Fig 1), Cyanobacteria is clearly the most abundant microflora in samples from amaranth grain varieties as shown in Fig 2. While Cyanobacteria is detectable in the soil around the 4 amaranth species studied here and is significantly abundant in rhizospheric samples, only in *A. hypochondriacus* and *A. cruentus* there is an explosion of Cyanobacteria in rhizospheric soil (AHRS and ARRS) and reduced representation from Proteobacteria and Actinobacteria. Although the abundance of Cyanobacteria in the rhizospheric soil in *A. caudatus* is more than the negative controls (non-amaranth plant species), it is still relatively lower than from the other two amaranth grain varieties. This sample also has a greater representation from Proteobacteria than the other two grain amaranth plants.

**Fig 1:**
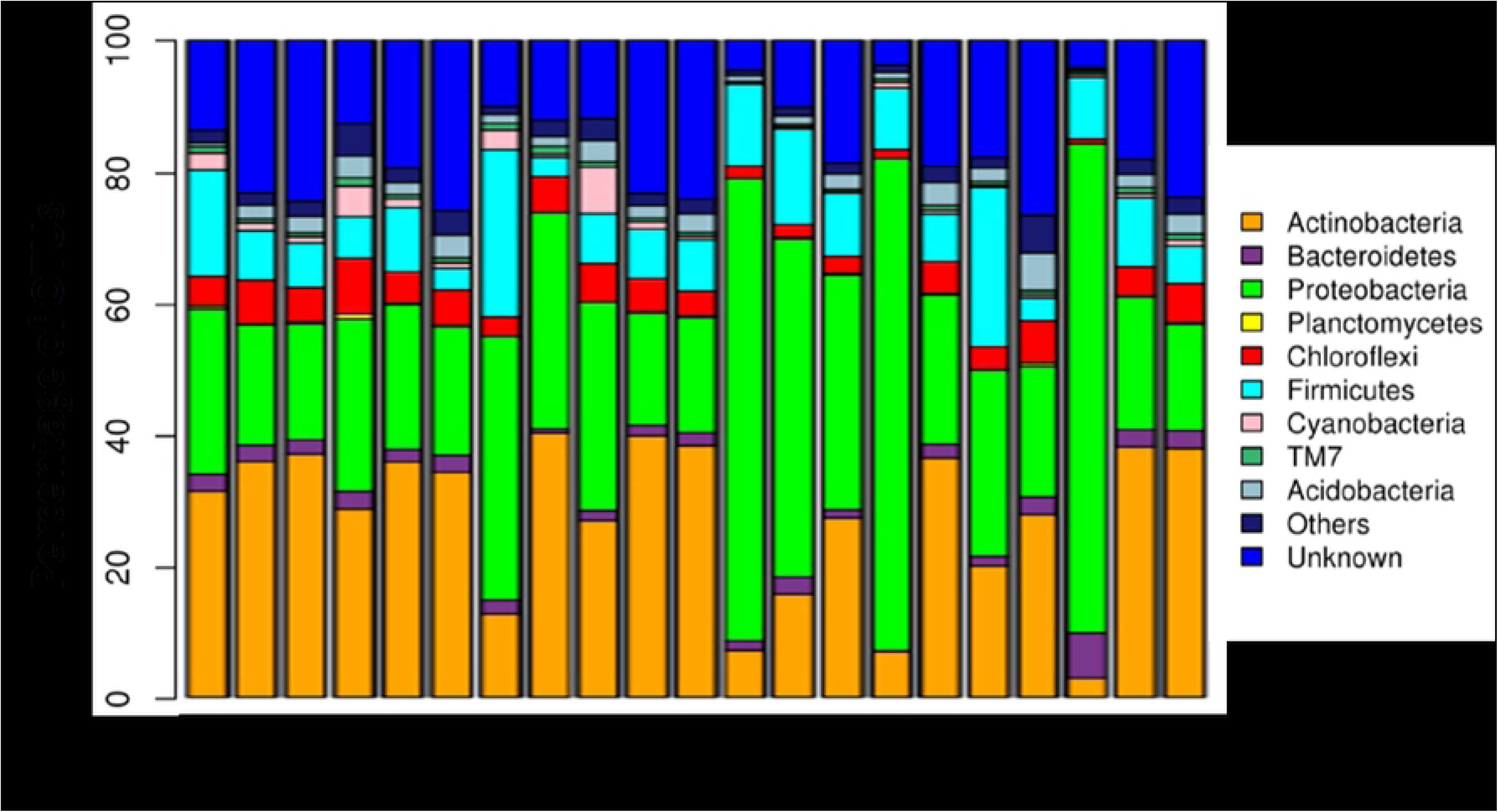
OTUs at phylum level. Top 10 phyla shown Sample key:AC -*A. caudatus;* AH -*A. hypochondriacus;* AR -*A. cruentus;* AM -*Amaranthus* wild type; BT -*Beta vulgaris*; CC -*Cicer arietinum*; TM -*Solanum lycopersicum*; SOIL -Bulk soil away from plants as control; R -Plant Root; RS -Rhizospheric soil; S – Outer rhizospheric soil. For eg., ACRS – *A. caudatus* (AC) rhizospheric soil (RS). The top 10 bacterial phyla from the QIIME pipeline analysis are represented as percentages of OTUs (y-axis) for all 21 samples (x-axis).

**Fig 2:**
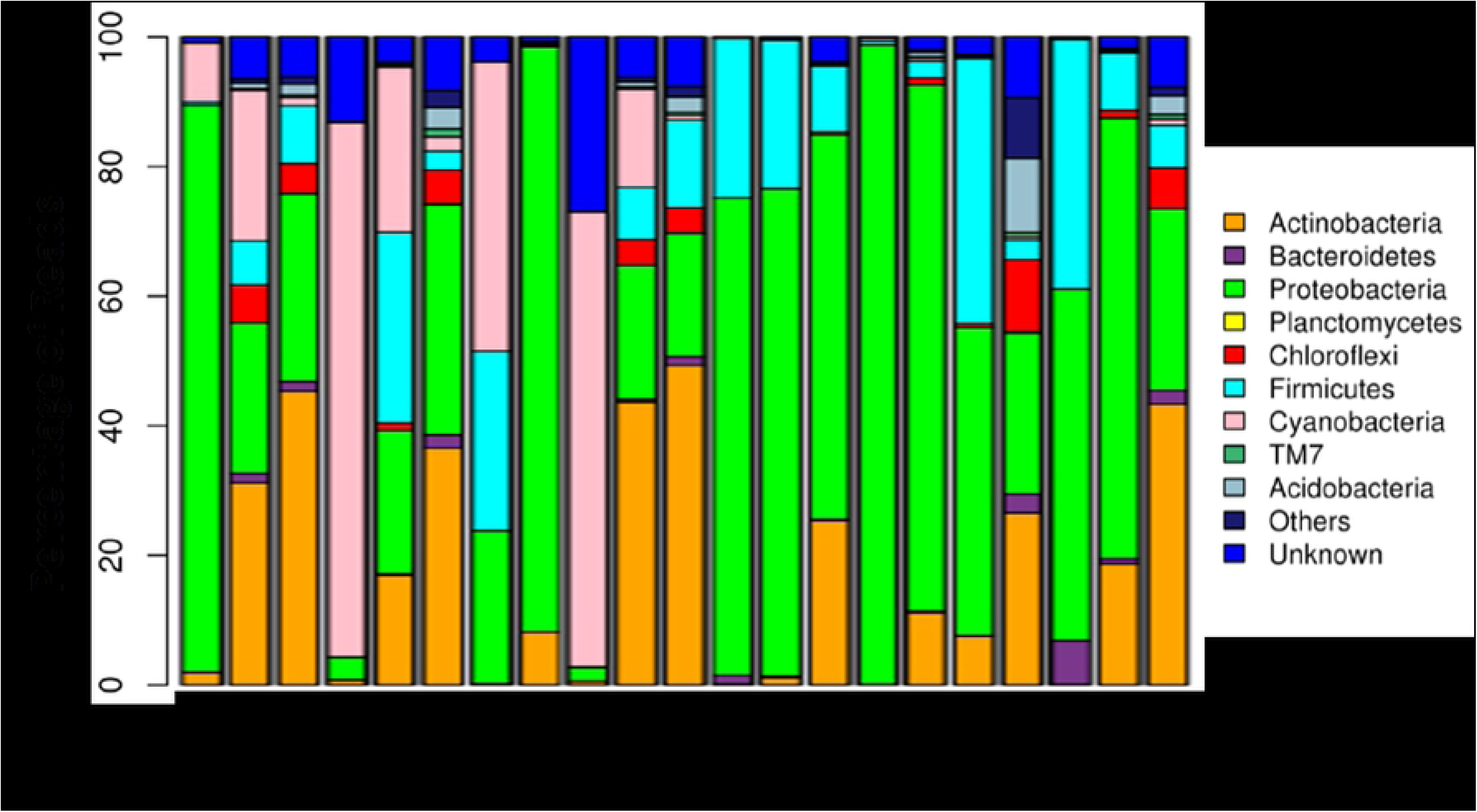
Reads at phylum level showing relative abundance. Top 10 phyla shown Sample key:AC -*A. caudatus;* AH -*A. hypochondriacus;* AR -*A. cruentus;* AM -*Amaranthus* wild type; BT -*Beta vulgaris*; CC -*Cicer arietinum*; TM -*Solanum lycopersicum*; SOIL -Bulk soil away from plants as control; R -Plant Root; RS -Rhizospheric soil; S – Outer rhizospheric soil. For eg., ACRS – *A. caudatus* (AC) rhizospheric soil (RS). The top 10 bacterial phyla from the QIIME pipeline analysis are represented in terms of percentages of Reads (y-axis) mapping to their respective OTUs for all 21 samples (x-axis).

### Microbial Diversity Analysis

Microbial diversity within the samples (alpha diversity) was measured by using metrics like Shannon, Chao1 and Observed species metrics. The Chao1 metric estimates the species richness while Shannon metric estimates observed OTU abundances and accounts for both richness and evenness. Shannon plots show that the rarefaction curves plateau after X **#** of reads which suggests that the depth of sequencing is sufficient to infer diversity in bacterial species. The Observed species metric is the count of unique OTUs identified in the sample. From the rarefaction curves, we observe that Bulk Soil sample has the highest species richness and evenness than other samples. It also has the highest count for unique OTUs as seen in Observed species metric. Most of the outer rhizospheric soil samples also show higher richness than the rhizospheric soil samples and root samples, which show lower OTU abundances. In root samples from *Amaranthus hypochondriacus, Amaranthus cruentus and Amaranthus caudatus* the richness was found to be the least compared to other plants and therefore abundance by one or a few kinds of OTUs seems to be prominent. Amaranth wild type showed much higher abundance of OTUs representing greater richness of the microbial communities in the roots. All other plants showed higher richness in diversity analyses (S1 Fig, S2 Fig and S3 Fig).

Beta diversity analysis was done using jackknifed test with max rarefaction limited to 250000 sequences per sample. The difference between the samples was explained by 3 component PCoA analysis (PC1 49.97%, PC2 22.93% and PC3 11.27%) (Fig 3). On the plot, most of the outer rhizospheric soil samples grouped together with bulk soil on the PC2 axis. *Amaranthus hypochondriacus* and *Amaranthus cruentus* root samples co-segregate on the PC3 axis and *Amaranthus caudatus* is located towards the PC1 axis suggesting more diversity of microbial community in the later than previous two. Amaranth rhizospheric soil samples appear to be distinct compared to other non-amaranth plant species.

**Fig 3:**
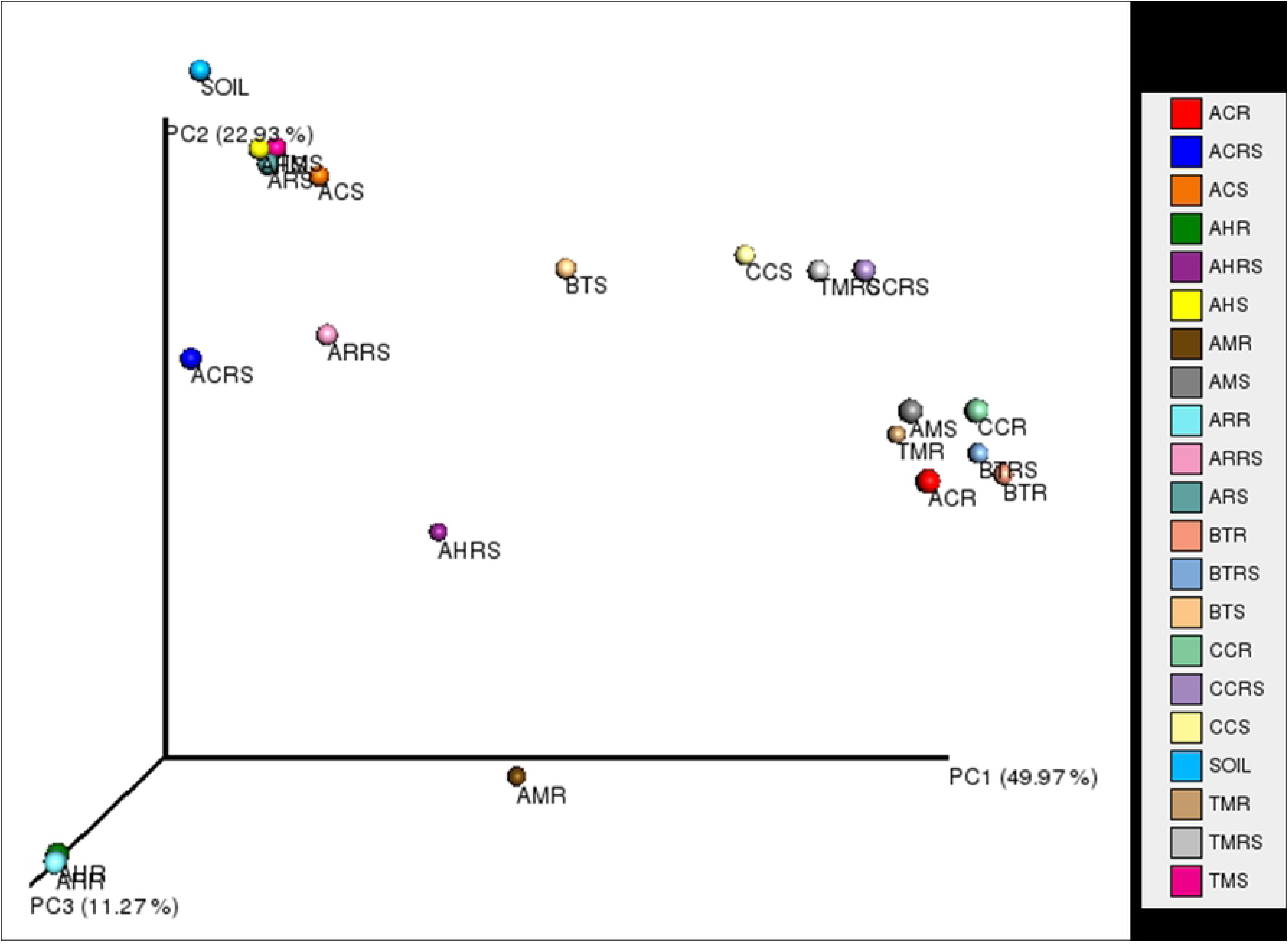
Beta diversity analysis: Principal Coordinate Analysis (PCoA) plot Sample key: AC -*A. caudatus;* AH -*A. hypochondriacus;* AR -*A. cruentus;* AM -*Amaranthus* wild type; BT -*Beta vulgaris*; CC -*Cicer arietinum*; TM -*Solanum lycopersicum*; SOIL -Bulk soil away from plants as control; R -Plant Root; RS -Rhizospheric soil; S – Outer rhizospheric soil. For eg., ACRS – *A. caudatus* (AC) rhizospheric soil (RS).

Beta diversity analysis using Jackknifed test was done with max rarefaction limited to 250000 sequences per sample. Principal Coordinate Analysis plot was generated from weighted UniFrac distance matrix for all 21 root, rhizosphere and soil samples and visualized using Emperor.

### OTU diversity in Cyanobacteria across samples

There are 3301 OTUs assigned to Cyanobacteria based on the criteria described in methods. The chart in Fig 4A shows the distribution of these OTU across samples. Relatively high numbers of OTUs are present in samples from amaranth species than other control species. A similar chart showing the distribution of OTUs from Proteobacteria (Fig 4B), which is an order of magnitude more than those of Cyanobacteria, show reduced number of OTUs in amaranth root and rhizosphere. This is despite the fact that rhizospheric soil from amaranth is rich in OTUs from Proteobacteria.

**Fig 4:**
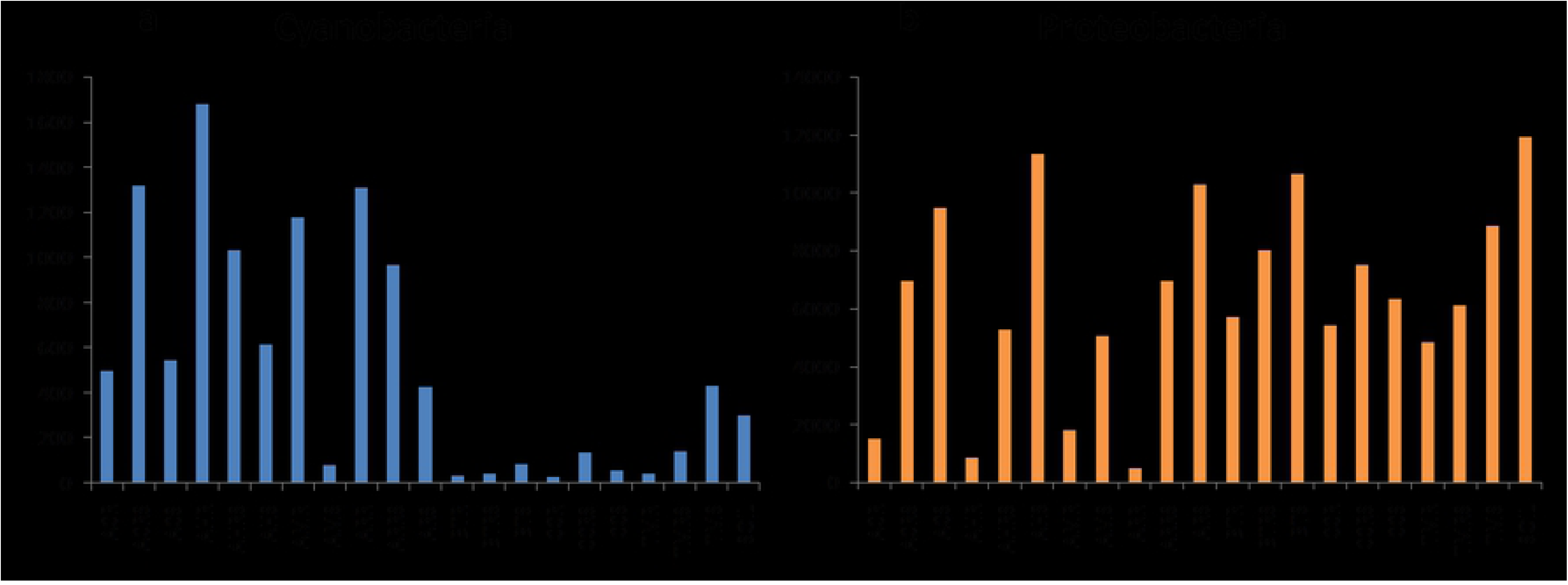
OTUs distribution in all samples Sample key: AC -*A. caudatus;* AH -*A. hypochondriacus;* AR -*A. cruentus;* AM -*Amaranthus* wild type; BT -*Beta vulgaris*; CC -*Cicer arietinum*; TM -*Solanum lycopersicum*; SOIL -Bulk soil away from plants as control; R -Plant Root; RS -Rhizospheric soil; S – Outer rhizospheric soil. For eg., ACRS – *A. caudatus* (AC) rhizospheric soil (RS). A: Cyanobacteria. B: Proteobacteria. Sample names are represented on x-axis and y-axis represents the number of OTUs from the samples without any cutoff for abundance.

### Alpha and beta diversity among Cyanobacteria in samples from Amaranth

Venn diagram (S4 Fig) of the highly abundant OTUs (1000 amplicons across all samples) from microbes with significant presence (minimum 30 amplicons per sample) suggest that compared to the alpha diversity in the three amaranth species, beta diversity is minimal. For examples, alpha diversity ranges from 17-18 OTUs in rhizospheric soil and ranges from 15-17 in root. However the alpha diversity varies in samples from outer rhizospheric soil, perhaps from the stringent cutoff of 30 amplicons per OTUs. In S4 Fig the case where when no cutoff is used for the OTUs beta diversity is observed in the *A. hypochondriacus* samples.

### Amaranth Root transcriptome analysis

A small number of hits were observed after mapping reads from root tissue of whole transcriptome analysis for *Amaranthus hypochondriacus* [42] onto bacterial genomes using Bowtie [46] aligner. Out of these, the number of hits from Proteobacteria was most abundant followed by Actinobacteria. But still it showed a pattern wherein the number of Cyanobacterial hits seen in the 30-day plant stage (AH12) were significantly higher than in 15-day plant stage (AH3) when compared to bacterial hits from other phyla (as seen in Fig 5).

**Fig 5:**
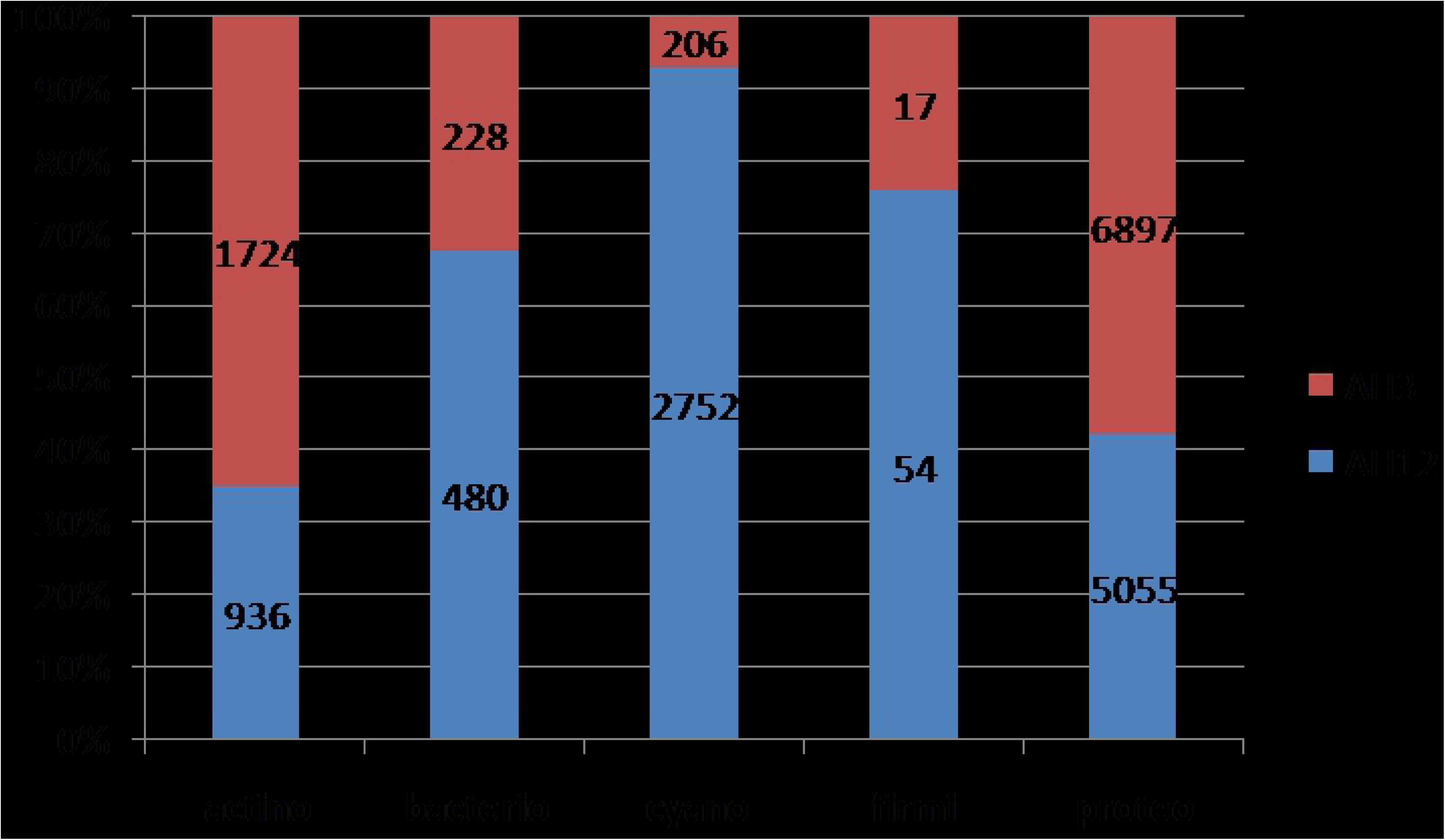
Major Phyla seen in Comparative Analysis of Amaranth Root transcriptomes for 15-day (AH3) and 30-day (AH12) stages Sample key: **actino** – Actinobacteria; **bacterio** – Bacteroidetes; **cyano** – Cyanobacteria; **firmi** – Firmicutes; **proteo** – Proteobacteria Bacterial hits observed from root transcriptome of *Amaranthus hypochondriacus* at 15-day stage (AH3) and 30-day stage (AH12) are shown in terms of percentages to reveal relative change in the bacterial population on Amaranth roots. The number of hits from the stages is represented on the histogram.

## DISCUSSION

Here, we report a comparative study of the root-associated microbiota between the three grain amaranths (viz. *A. hypochondriacus, A.cruentus, A. caudatus*) and three other plant species under different order (viz. *Beta vulgaris, Cicer arietinum* and *Solanum lycopersicum*) as negative control. We employed 16S rRNA amplicon sequencing for the purpose of profiling microbial species associated with the roots of these plants to find the microflora specifically being enriched in grain amaranths. We observed that Cyanobacteria are distinctly enriched in the roots of grain amaranths in terms of abundance and uniqueness. Though most dominant phyla are Proteobacteria and Actinobacteria, the most unique distinct feature of phyla level of analysis is the presence of Cyanobacterial otus in amaranth roots, *A. hypochondriacus and A. cruentus*. *A. caudatus* was different from the other two in that it shows higher number of Proteobacteria and lower number of Cyanobacteria. This could be due to the differences in the plant genotypes as *A. caudatus* plants grow better under higher altitudes compared to *A. cruentus* and *A. hypochondriacus.*

Both photosynthetic and non-photosynthetic Cyanobacteria have been known to associate with plants for carbon source and some of these have ability to fix nitrogen which is beneficial for plants [47]. Although this is not known for grain Amaranths which unlike legumes like *Cicer* do not form root nodules, it might be presumed that the enriched bacteria might be helping in the fitness of the plant for survival. Proteobacteria was seen enriched beet root and *Cicer* plants (Fig 2) which could also be playing a role in plant protection [48].

Microbial diversity analyses reveal that the microbial community profiles of *Amaranthus hypochondriacus* and *Amaranthus cruentus* root samples and rhizosphere samples were distinct in composition compared to the other non amaranth species described in the study (Fig 3). One of the prime reasons is the composition of OTUs coming from Cyanobacteria.

Abundance of 16S rRNA under Cyanobacteria needs to be reconciled from comparison with Amaranth chloroplast. Firstly, Cyanobacterial OTUs are more diverse than is expected from chloroplast of *A. hypochondriacus*. Secondly, the extraction procedure and sequencing methodology being the same for all species studied here, the enrichment observed in Cyanobacterial OTUs for amaranth plants was significantly greater than for non-amaranth plants. Thirdly, the tissue in question in this study is root, which unlike leaves or shoot, are not known to have chloroplasts. Additionally we also observed enrichment of Cyanobacteria derived transcripts in amaranth developmental transcriptome, when the 15-day and 30-day *A. hypochondriacus* plants were compared. Lastly, to check whether we can find culturable Cyanobacteria associated with Amaranth plants we isolated cyanobacterial species from *A. hypochondriacus* roots resembling at least four different species (data unpublished).

Earlier studies have shown that cyanobacteria can form associations in general with plants like cycads and *Gunnera* and can fix nitrogen in soil either as free-living organisms or in association with host plants where they reside in specific tissues of the host. There are also many examples which bring to light the utility of cyanobacteria like *Calothrix* and *Anabena* which can be used as biofertilizers in conjunction with other rhizobacteria or alone and can aid in the nutritional profile and fitness of plants [49, 50, 51, 52, 53, 54].

Considering that grain species under Amaranthaceae skipped the genome duplication event during the Oligocene period common to plant families under other orders perhaps as a survival mechanism [42] and considering that it has amassed unusually high numbers of desirable traits, it is tempting to hypothesize that species under the family Amarathaceae may have used alternate mechanism to escape the stress during the Oligocene period, which may include symbiosis with unique microbes. Even though the role of cyanobacteria in Amaranth plants needs further investigation, it is possible that cyanobacteria are recruited by Amaranth plants from soil and that they play a helpful role favouring the fitness of Amaranth plants and perhaps even improve their nutritional profile. The associated microbes could also be investigated for their future potential to transfer useful properties to other crop species under cultivation by plant microbiome engineering [55,56].

## MATERIALS AND METHODS

### Sample collection and preparation

Six plant species were selected for 16S rRNA metagenomic profiling including *Amaranthus hypochondriacus, Amaranthus cruentus, Amaranthus caudatus, Cicer arietinum, Solanum lycopersicum*, and *Beta vulgaris*. The plants were grown in-house on tilled plots (Karnataka red soil) at IBAB, Bangalore where the average day time temperatures vary from 15-30 °C. All plants were 30 to 35 days old when harvested. Whole plants were extracted along with the soil attached to the roots, flash frozen in liquid nitrogen and stored in −80 °C freezer until processed for metagenomic DNA isolation. Apart from this, soil from the same plot was collected separately as Bulk soil and stored as mentioned above. In addition to this, wild Amaranth plants growing in the campus were also incorporated for comparison.

Triplicates of plant root samples (biological replicates) were collected and metagenomic DNA isolation for the same was done and pooled as a single sample for this work. For processing the samples for metagenomic isolation, three fractions were collected from each sample. The plant roots were shaken vigorously to detach loosely bound soil which was collected as outer rhizospheric soil. The soil that remained adhered to the plant surface was gently scraped off and collected as rhizospheric soil and then the roots were washed thoroughly with distilled water. The washed roots were homogenized using pestle and mortar and collected as root extract.

For root transcriptome sequencing, 15 and 30 days old *Amaranthus hypochondriacus* plants were extracted from soil, root tissue was excised and cleaned with RNase-free water and bleach (4% for 10 mins.), then washed thoroughly in RNase-free water, flash frozen in liquid nitrogen and stored in −80 °C freezer until RNA extraction was done (42).

### Isolation of Metagenomic DNA

Metagenomic DNA (Mg-DNA) was isolated from the samples using the kit FastDNA™ SPIN Kit for Soil (MP Biomedicals). In brief, 0.3 to 0.5 g of sample was added to each lysing matrix E bead tube and vortexed for maximum 10 minutes for soil lysis in buffer. This was proceeded with Mg-DNA isolation as per kit instructions. Final elution was done in 100 µL of kit supplied RNase-free DES water after incubation at 55°C for 5 min. Purity and concentration of eluted DNA was checked using Nanodrop 1000 spectrophotometer. Isolation of Mg-DNA was confirmed using full length 16S rRNA PCR amplification. The primers used for the PCR were: Forward primer sequence −27F: 5’ AGAGTTTGATCMTGGCTCAG 3’ and Reverse primer sequence −1492R: 5’ TACGGYTACCTTGTTACGACTT 3’. 20 μL of PCR reaction mix was made up with 0.2 μL template (∼ 30 −40 ng), 1X KAPA Taq Buffer A, 2.5 mM MgCl_2_, 2.0 U KAPA Taq DNA polymerase (KAPA Biosystems), 1 mM of dNTPs, 0.7 μM of each primer and Milli-Q sterile water. The PCR conditions employed for full length 16SrRNA PCR amplification include i) initial denaturation for 5 min at 95°C, ii) followed by 30 cycles of (45 s, 94°C; 1 min, 54°C; 1.5 min, 72°C), and iii) final extension of 5 min at 72°C. Amplified samples were visualized using agarose gel electrophoresis (1%) stained with ethidium bromide and run alongside 1 μL of 2-log DNA ladder.

### Isolation of root total RNA

Root tissues collected from 15-day and 30-day plants and stored in −80°C freezer were homogenized using RNAse-free pestle and mortar. Total RNA from root tissues were extracted using standardized Trizol extraction method. Samples for root tissue from 15 days old plants were pooled together during homogenization step due to low amount of tissue available [42].

### 16S rRNA Library Preparation and 16SrRNA Amplicon Sequencing

In total 21 DNA samples were sent for library preparation and sequencing at SciGenom Labs Pvt Ltd, Kochi. PCR amplification of V3 region was done using PCR forward primer sequence- 341 Forward: 5’ CCTACGGGAGGCAGCAG 3’ and reverse primer sequence- 518 Reverse: 5’ ATTACCGCGGCTGCTGG 3’. The PCR conditions were as follows: PCR Master Mix contained 2 μL each 10 pmol/µL forward and reverse primers, 0.5 μL of 40mM dNTP, 5 μL of 5X Phusion HF reaction buffer, 0.2 μL of 2U/µL F-540 Special Phusion HS DNA Polymerase, 5ng input DNA and water to make up the total volume to 25 μL. The PCR reaction was 98°C for 30 sec; 30 cycles of 98°C for 10 sec, 72°C for 30 sec followed by extension at 72°C for 5 sec and 4°C hold. Size selection was done using gel extraction of 230 to 250 bp region. Primers with proprietary barcodes were used for the sequencing experiment and the Illumina adapter and index sequences were attached to the size-selected V3 amplicons using the following PCR conditions: PCR Master Mix will contain 2 μL each 10 pmol/µL forward and reverse primers, 1 μL of 40mM dNTP, 10 μL of 5 X Phusion HF reaction buffers, 0.4 μL of 2U/µL F-540 Special Phusion HS DNA Polymerase, 10 μL (minimum 5 ng) of PCR amplicon and water to make up the total volume to 50 μL. The PCR reaction was 98°C for 30 sec; 15 cycles of 98°C for 10 sec, 72°C for 30 sec followed by extension at 72°C for 5 sec and 4°C hold. Size selection was done using gel extraction of 354 to 362 bp region. Multiplexed paired-end amplicon sequencing for V3 region of 16S rRNA was done using Illumina MiSeq platform for all 21 DNA samples at a depth of about 0.5 million paired-end reads per sample and a read length of 150 bases.

### RNA library preparation and sequencing

One microgram of the total RNA extracted from the same type of tissue but from the three different batches of the plants were pooled together to make 3 μg of the starting material for library preparation as measured by the NanoDrop Spectrophotometer. Further, transcriptome libraries were prepared using TruSeq RNA Sample Preparation Low Throughput (LT) protocol (Illumina) as per the manufacturer’s guidelines and as reported previously [42]. The quality and quantity of the libraries were estimated by Qubit fluorometry (Invitrogen), and the size distribution was analysed on Bioanalyzer (Agilent) using high sensitivity DNA chips.

### Data Analysis

Raw sequenced reads were trimmed for spacer and other unwanted sequences and were passed through FastQC [43] reads filter. The good quality reads were merged using PANDAseq [44]. The assembled reads were used for downstream analysis using QIIME pipeline (v1.9.1) [45]. Operational taxonomic units (OTUs) were clustered based on sequence similarity at 97% using UCLUST [57]. Representative sequences were identified and aligned against Greengenes [58] core set of sequences using PyNAST [59]. The chimeras were removed using UCHIME [60] and the filtered alignment was used for building phylogenetic tree using FastTree [61]. OTU table was prepared from predicted OTUs and UCLUST assigned taxonomy file from which singletons were removed from analysis. Alpha diversity rarefaction curves were used to measure microbial diversity within the samples by calculating Shannon, Chao1 and Observed Species metrics using default parameters and maximum rarefaction depth of 250000. For Beta diversity analysis a jackknifed test was performed to construct a consensus UPGMA tree using weighted UniFrac [62] distance matrix and a PCoA plot was generated for all 21 samples and visualized using Emperor [63].

### Mining Amaranth Root transcriptome for bacterial transcripts

Root transcriptomes of 15-day stage and 30-day stage of *Amaranthus hypochondriacus* [40, 42], one of the grain amaranths were mined for the presence of bacterial transcripts and the bacterial hits observed were compared between the two stages. The transcriptomic reads were mapped using Bowtie [46] in try-hard mode onto microbial genomes (bacteria and archaea) downloaded from NCBI.

## ACKNOWLEDGEMENTS

The authors acknowledge Department of Biotechnology, Government of India (GoI) for BINC fellowship to Suran Nambisan, Ramalingaswamy fellowship from DBT to Subhashini Srinivasan (BT/HRD/35/02/17/2009) and DBT grants to both Bibha Choudhary and Subhashini Srinivasan (BTPR12422/MED/31/287/2014, valid from November 2014 to 2017). The authors wish to acknowledge DST-FIST, Government of India and Department of IT, BT and S&T, Government of Karnataka for computing infrastructure to IBAB. The authors also acknowledge SciGenom Pvt. Ltd., Kochi for 16S rRNA amplicon sequencing done for this study.

## AUTHOR CONTRIBUTIONS

**Conceptualization:** Bibha Choudhary, Subhashini Srinivasan

**Data curation:** Suran Nambisan, Meeta Sunil

**Formal analysis:** Suran Nambisan

**Funding acquisition:** Bibha Choudhary, Subhashini Srinivasan.

**Investigation:** Suran Nambisan, Meeta Sunil, Bibha Choudhary, Subhashini Srinivasan

**Methodology:** Suran Nambisan, Meeta Sunil, Bibha Choudhary, Subhashini Srinivasan

**Project administration:** Suran Nambisan, Bibha Choudhary, Subhashini Srinivasan

**Resources:** Bibha Choudhary, Subhashini Srinivasan

**Software:** Suran Nambisan, Subhashini Srinivasan

**Supervision:** Bibha Choudhary, Subhashini Srinivasan

**Validation:** Suran Nambisan, Bibha Choudhary, Subhashini Srinivasan

**Visualization:** Suran Nambisan, Subhashini Srinivasan

**Writing - original draft:** Suran Nambisan, Subhashini Srinivasan

**Writing - review & editing:** Suran Nambisan, Bibha Choudhary, Subhashini Srinivasan

## DATA AVAILABILITY

Raw reads from 16S rRNA sequence from 21 samples is available in the NCBI SRA database under the accession number SRP124254 under the NCBI Bioproject PRJNA417229 and root transcriptome from two time points (15 and 30 days) is available under NCBI accession number SRP111547.

## SUPPORTING INFORMATION

**S1 Fig.** Alpha diversity analysis: Shannon rarefaction curve for all 21 samples (TIF)

**S2 Fig.** Alpha diversity analysis: Chao1 rarefaction curve for all 21 samples (TIF)

**S3 Fig.** Alpha diversity analysis: Observed species rarefaction curve for all 21 samples (TIF)

**S4 Fig.** Venn Diagrams for Cyanobacterial OTUs in Amaranth samples (TIF)

**S1 Table.** Summary of All Reads from 16S rRNA V3 Amplicon Sequencing Experiment (XLSX)

## REFERENCES

1. Philippot L, Raaijmakers JM, Lemanceau P, van der Putten WH. Going back to the roots: the microbial ecology of the rhizosphere. Nat Rev Microbiol. 2013 Nov;11(11):789–99. doi: 10.1038/nrmicro3109. PMID: 24056930

2. Bulgarelli D, Schlaeppi K, Spaepen S, van Themaat EVL, Schulze-Lefert P. Structure and functions of the bacterial microbiota of plants. Annu Rev Plant Biol. 2013 Apr 29;64(1):807–38. doi: 10.1146/annurev-arplant-050312-120106. PMID: 23373698

3. Chaparro JM, Badri DV, Vivanco JM. Rhizosphere microbiome assemblage is affected by plant development. ISME J. 2014 Apr;8(4):790–803. doi: 10.1038/ismej.2013.196. PMID: 24196324

4. Chardot-Jacques V, Calvaruso C, Simon B, Turpault M-P, Echevarria G, Morel J-L. Chrysotile dissolution in the rhizosphere of the nickel hyperaccumulator *Leptoplax emarginata*. Environ Sci Technol. 2013 Mar 19;47(6):2612–20. doi: 10.1021/es301229m. PMID: 23373689

5. Ravikumar S, Ignatiammal STM, Gnanadesigan M, Kalaiarasi A. Effects of saline tolerant Azospirillum species on the growth parameters of mangrove seedlings. J Environ Biol. 2012 Sep;33(5):933–9. PMID: 23734462

6. Maróti G, Kondorosi É. Nitrogen-fixing *Rhizobium*-legume symbiosis: are polyploidy and host peptide-governed symbiont differentiation general principles of endosymbiosis? Front Microbiol. 2014 Jun 30;5:326. doi: 10.3389/fmicb.2014.00326. PMID: 25071739

7. Kunert KJ, Vorster BJ, Fenta BA, Kibido T, Dionisio G, Foyer CH. Drought stress responses in soybean roots and nodules. Front Plant Sci. 2016 Jul 12;7:1015. doi: 10.3389/fpls.2016.01015. PMID: 27462339

8. Harrison MJ. Signaling in the arbuscular mycorrhizal symbiosis. Annu Rev Microbiol. 2005 Oct;59(1):19–42. doi: 10.1146/annurev.micro.58.030603.123749. PMID: 16153162

9. Chen C, Zhu H. Are common symbiosis genes required for endophytic rice-rhizobial interactions? Plant Signal Behav. 2013 Sep;8(9). pii: e25453. doi: 10.4161/psb.25453. PMID: 23838959

10. Mousa WK, Shearer C, Limay-Rios V, Ettinger CL, Eisen JA, Raizada MN. Root-hair endophyte stacking in finger millet creates a physicochemical barrier to trap the fungal pathogen *Fusarium graminearum*. Nat Microbiol. 2016 Sep 26;1:16167. doi: 10.1038/nmicrobiol.2016.167. PMID: 27669453

11. de Vrieze J. The littlest farmhands. Science. 2015 Aug 14;349(6249):680–3. doi: 10.1126/science.349.6249.680. PMID: 26273035

12. Mayak S, Tirosh T, Glick BR. Plant growth-promoting bacteria confer resistance in tomato plants to salt stress. Plant Physiol Biochem. 2004 Jun;42(6):565–72. doi: 10.1016/j.plaphy.2004.05.009. PMID: 15246071

13. Pieterse CMJ, Zamioudis C, Berendsen RL, Weller DM, Van Wees SCM, Bakker PAHM. Induced systemic resistance by beneficial microbes. Annu Rev Phytopathol. 2014 Aug 4;52(1):347–75. doi: 10.1146/annurev-phyto-082712-102340. PMID: 24906124

14. Kalam S, Das SN, Basu A, Podile AR. Population densities of indigenous *Acidobacteria* change in the presence of plant growth promoting rhizobacteria (PGPR) in rhizosphere. J Basic Microbiol. 2017 May;57(5):376–385. doi: 10.1002/jobm.201600588. PMID: 28397264

15. Zamioudis C, Pieterse CMJ. Modulation of host immunity by beneficial microbes. Mol Plant Microbe Interact. 2012 Feb;25(2):139–50. doi: 10.1094/MPMI-06-11-0179. PMID: 21995763

16. Schenk PM, Carvalhais LC, Kazan K. Unraveling plant–microbe interactions: can multi-species transcriptomics help? Trends Biotechnol. 2012 Mar;30(3):177–84. doi: 10.1016/j.tibtech.2011.11.002. PMID: 22209623

17. Schloissnig S, Arumugam M, Sunagawa S, Mitreva M, Tap J, Zhu A, et al. Genomic variation landscape of the human gut microbiome. Nature. 2013 Jan 3;493(7430):45–50. doi: 10.1038/nature11711. PMID: 23222524

18. Keshri J, Mishra A, Jha B. Microbial population index and community structure in saline–alkaline soil using gene targeted metagenomics. Microbiol Res. 2013 Mar 30;168(3):165–73. doi: 10.1016/j.micres.2012.09.005. PMID: 23083746

19. Albertsen M, Karst SM, Ziegler AS, Kirkegaard RH, Nielsen PH. Back to basics – the influence of DNA extraction and primer choice on phylogenetic analysis of activated sludge communities. PLoS One. 2015 Jul 16;10(7):e0132783. doi: 10.1371/journal.pone.0132783. PMID: 26182345

20. Håvelsrud O, Haverkamp TH, Kristensen T, Jakobsen KS, Rike A. Metagenomic and geochemical characterization of pockmarked sediments overlaying the Troll petroleum reservoir in the North Sea. BMC Microbiol. 2012 Sep 11;12:203. doi: 10.1186/1471-2180-12-203. PMID: 22966776

21. Bulgarelli D, Rott M, Schlaeppi K, Ver Loren van Themaat E, Ahmadinejad N, Assenza F, et al. Revealing structure and assembly cues for *Arabidopsis* root-inhabiting bacterial microbiota. Nature. 2012 Aug 2;488(7409):91–5. doi: 10.1038/nature11336. PMID: 22859207

22. Lundberg DS, Lebeis SL, Paredes SH, Yourstone S, Gehring J, Malfatti S, et al. Defining the core *Arabidopsis thaliana* root microbiome. Nature. 2012 Aug 2;488(7409):86–90. doi: 10.1038/nature11237. PMID: 22859206

23. Bodenhausen N, Horton MW, Bergelson J. Bacterial communities associated with the leaves and the roots of *Arabidopsis thaliana*. PLoS One. 2013;8(2):e56329. doi: 10.1371/journal.pone.0056329. PMID: 23457551

24. Haney CH, Samuel BS, Bush J, Ausubel FM. Associations with rhizosphere bacteria can confer an adaptive advantage to plants. Nat Plants. 2015 May 11;1(6):15051. doi: 10.1038/nplants.2015.51. PMID: 27019743

25. Bai Y, Müller DB, Srinivas G, Garrido-Oter R, Potthoff E, Rott M, et al. Functional overlap of the *Arabidopsis* leaf and root microbiota. Nature. 2015 Dec 17;528(7582):364–9. doi: 10.1038/nature16192. PMID: 26633631

26. Zgadzaj R, Garrido-Oter R, Jensen DB, Koprivova A, Schulze-Lefert P, Radutoiu S. Root nodule symbiosis in *Lotus japonicus* drives the establishment of distinctive rhizosphere, root, and nodule bacterial communities. Proc Natl Acad Sci. 2016 Dec 6;113(49):E7996–E8005. doi: 10.1073/pnas.1616564113. PMID: 27864511

27. Camps C, Jardinaud M-F, Rengel D, Carrère S, Hervé C, Debellé F, et al. Combined genetic and transcriptomic analysis reveals three major signalling pathways activated by Myc-LCOs in *Medicago truncatula*. New Phytol. 2015 Oct;208(1):224–40. doi: 10.1111/nph.13427. PMID: 25919491

28. Rascovan N, Carbonetto B, Perrig D, Díaz M, Canciani W, Abalo M, et al. Integrated analysis of root microbiomes of soybean and wheat from agricultural fields. Sci Rep. 2016 Jun 17;6:28084. doi: 10.1038/srep28084. PMID: 27312589

29. Edwards J, Johnson C, Santos-Medellín C, Lurie E, Podishetty NK, Bhatnagar S, et al. Structure, variation, and assembly of the root-associated microbiomes of rice. Proc Natl Acad Sci. 2015 Feb 24;112(8):E911–20. doi: 10.1073/pnas.1414592112. PMID: 25605935

30. Breidenbach B, Pump J, Dumont MG. Microbial community structure in the rhizosphere of rice plants. Front Microbiol. 2016 Jan 13;6:1537. doi: 10.3389/fmicb.2015.01537. PMID: 26793175

31. Peiffer JA, Spor A, Koren O, Jin Z, Tringe SG, Dangl JL, et al. Diversity and heritability of the maize rhizosphere microbiome under field conditions. Proc Natl Acad Sci. 2013 2013 Apr 16;110(16):6548–53. doi: 10.1073/pnas.1302837110. PMID: 23576752

32. Correa-Galeote D, Bedmar EJ, Fernández-González AJ, Fernández-López M, Arone GJ. Bacterial communities in the rhizosphere of amilaceous maize (*Zea mays* L.) as assessed by pyrosequencing. Front Plant Sci. 2016 Jul 29;7:1016. doi: 10.3389/fpls.2016.01016. PMID: 27524985

33. Bulgarelli D, Garrido-Oter R, Münch PC, Weiman A, Dröge J, Pan Y, et al. Structure and function of the bacterial root microbiota in wild and domesticated barley. Cell Host Microbe. 2015 Mar 11;17(3):392–403. doi: 10.1016/j.chom.2015.01.011. PMID: 25732064

34. Zachow C, Müller H, Tilcher R, Berg G. Differences between the rhizosphere microbiome of Beta vulgaris ssp. maritima - ancestor of all beet crops - and modern sugar beets. Front Microbiol. 2014 Aug 26;5:415. doi: 10.3389/fmicb.2014.00415. PMID: 25206350

35. Shi Y, Yang H, Zhang T, Sun J, Lou K. Illumina-based analysis of endophytic bacterial diversity and space-time dynamics in sugar beet on the north slope of Tianshan mountain. Appl Microbiol Biotechnol. 2014;98(14):6375–85. doi: 10.1007/s00253-014-5720-9. PMID: 24752839

36. Lee SA, Park J, Chu B, Kim JM, Joa J-H, Sang MK, et al. Comparative analysis of bacterial diversity in the rhizosphere of tomato by culture-dependent and -independent approaches. J Microbiol. 2016 Dec;54(12):823–831. PMID: 27888459 doi: 10.1007/s12275-016-6410-3

37. İnceoğlu Ö, Al-Soud WA, Salles JF, Semenov AV, van Elsas JD. Comparative analysis of bacterial communities in a potato field as determined by pyrosequencing. Gilbert JA, editor. PLoS One. 2011;6(8):e23321. doi: 10.1371/journal.pone.0023321. PMID: 21886785

38. Wagner MR, Lundberg DS, del Rio TG, Tringe SG, Dangl JL, Mitchell-Olds T. Host genotype and age shape the leaf and root microbiomes of a wild perennial plant. Nat Commun. 2016 Jul 12;7:12151. doi: 10.1038/ncomms12151. PMID: 27402057

39. Shakya M, Gottel N, Castro H, Yang ZK, Gunter L, Labbé J, et al. A multifactor analysis of fungal and bacterial community structure in the root microbiome of mature *Populus deltoides* trees. PLoS One. 2013 Oct 16;8(10):e76382. doi: 10.1371/journal.pone.0076382. PMID: 24146861

40. Sunil M, Hariharan AK, Nayak S, Gupta S, Nambisan SR, Gupta RP, et al. The draft genome and transcriptome of *Amaranthus hypochondriacus*: a C4 dicot producing high-lysine edible pseudo-cereal. DNA Res. 2014 Dec;21(6):585–602. doi: 10.1093/dnares/dsu021. PMID: 25071079

41. Clouse JW, Adhikary D, Page JT, Ramaraj T, Deyholos MK, Udall JA, et al. The Amaranth genome: genome, transcriptome, and physical map assembly. Plant Genome. 2016 Mar;9(1):1–14. doi: 10.3835/plantgenome2015.07.0062. PMID: 27898770.

42. Sunil M, Hariharan N, Dixit S, Choudhary B, Srinivasan S. Differential genomic arrangements in Caryophyllales through deep transcriptome sequencing of *A. hypochondriacus*. PLoS One. 2017 Aug 7;12(8):e0180528. doi: 10.1371/journal.pone.0180528. PMID: 28786999

43. Andrews S. FastQC: a quality control tool for high throughput sequence data. 2010. Available from: https://www.bioinformatics.babraham.ac.uk/projects/fastqc/.

44. Masella AP, Bartram AK, Truszkowski JM, Brown DG, Neufeld JD. PANDAseq: paired-end assembler for illumina sequences. BMC Bioinformatics. 2012 Feb 14;13:31. doi: 10.1186/1471-2105-13-31. PMID: 22333067

45. Caporaso JG, Kuczynski J, Stombaugh J, Bittinger K, Bushman FD, Costello EK, et al. QIIME allows analysis of high-throughput community sequencing data. Nat Methods. 2010 May;7(5):335–6. doi: 10.1038/nmeth.f.303. PMID: 20383131

46. Langmead B, Trapnell C, Pop M, Salzberg SL. Ultrafast and memory-efficient alignment of short DNA sequences to the human genome. Genome Biol. 2009;10(3):R25. doi: 10.1186/gb-2009-10-3-r25. PMID: 19261174

47. Rai AN, Söderbäck E, Bergman B. Tansley Review No. 116. Cyanobacterium–plant symbioses. New Phytol. 2000;(147):449–481. doi: 10.1046/j.1469-8137.2000.00720.x

48. Mendes R, Kruijt M, de Bruijn, Dekkers E, van der Voort M, Sneider JH et al. Deciphering the rhizosphere microbiome for disease-suppressive bacteria. Science. 2011;332(6033):1097–100; doi: 10.1126/science.1203980. PMID: 21551032

49. Bidyarani N, Prasanna R, Chawla G, Babu S, Singh R. Deciphering the factors associated with the colonization of rice plants by cyanobacteria. J Basic Microbiol. 2015 Apr;55(4):407–19. doi: 10.1002/jobm.201400591. PMID: 25515189

50. Priya H, Prasanna R, Ramakrishnan B, Bidyarani N, Babu S, Thapa S, et al. Influence of cyanobacterial inoculation on the culturable microbiome and growth of rice. Microbiol Res. 2015 Feb;171:78–89. doi: 10.1016/j.micres.2014.12.011. PMID: 25644956

51. Manjunath M, Kanchan A, Ranjan K, Venkatachalam S, Prasanna R, Ramakrishnan B, et al. Beneficial cyanobacteria and eubacteria synergistically enhance bioavailability of soil nutrients and yield of okra. Heliyon. 2016 Feb 8;2(2):e00066. doi: 10.1016/j.heliyon.2016.e00066. PMID: 27441245

52. Prasanna R, Joshi M, Rana A, Shivay YS, Nain L. Influence of co-inoculation of bacteria-cyanobacteria on crop yield and C–N sequestration in soil under rice crop. World J Microbiol Biotechnol. 2012 Mar;28(3):1223–35. doi: 10.1007/s11274-011-0926-9. PMID: 22805842

53. Bidyarani N, Prasanna R, Babu S, Hossain F, Saxena AK. Enhancement of plant growth and yields in chickpea (*Cicer arietinum* L.) through novel cyanobacterial and biofilmed inoculants. Microbiol Res. 2016 Jul-Aug;188-189:97–105. doi: 10.1016/j.micres.2016.04.005. PMID: 27296967

54. Singh DP, Prabha R, Yandigeri MS, Arora DK. Cyanobacteria-mediated phenylpropanoids and phytohormones in rice (*Oryza sativa*) enhance plant growth and stress tolerance. Antonie Van Leeuwenhoek. 2011 Nov;100(4):557–68. doi: 10.1007/s10482-011-9611-0. PMID: 21732035

55. Coleman-Derr D, Tringe SG. Building the crops of tomorrow: advantages of symbiont-based approaches to improving abiotic stress tolerance. Front Microbiol. 2014 Jun 6;5:283. doi: 10.3389/fmicb.2014.00283. PMID: 24936202

56. Hu J, Wei Z, Friman V-P, Gu S, Wang X, Eisenhauer N, et al. Probiotic diversity enhances rhizosphere microbiome function and plant disease suppression. mBio. 2016 Dec 13;7(6). pii: e01790-16. doi: 10.1128/mBio.01790-16. PMID: 27965449

57. Edgar RC. Search and clustering orders of magnitude faster than BLAST. Bioinformatics. 2010 Oct 1;26(19):2460–1. doi: 10.1093/bioinformatics/btq461. PMID: 20709691

58. DeSantis TZ, Hugenholtz P, Larsen N, Rojas M, Brodie EL, Keller K, et al. Greengenes, a chimera-checked 16S rRNA gene database and workbench compatible with ARB. Appl Environ Microbiol. 2006 Jul;72(7):5069–72. doi: 10.1128/AEM.03006-05. PMID: 16820507

59. Caporaso JG, Bittinger K, Bushman FD, DeSantis TZ, Andersen GL, Knight R. PyNAST: a flexible tool for aligning sequences to a template alignment. Bioinformatics. 2010 Jan 15;26(2):266–7. doi: 10.1093/bioinformatics/btp636. PMID: 19914921

60. Edgar RC, Haas BJ, Clemente JC, Quince C, Knight R. UCHIME improves sensitivity and speed of chimera detection. Bioinformatics. 2011 Aug 15;27(16):2194–200. doi: 10.1093/bioinformatics/btr381. PMID: 21700674

61. Price MN, Dehal PS, Arkin AP. FastTree 2 – approximately maximum-likelihood trees for large alignments. PLoS One. 2010 Mar 10;5(3):e9490. doi: 10.1371/journal.pone.0009490. PMID: 20224823

62. Lozupone C, Knight R. UniFrac: a new phylogenetic method for comparing microbial communities. Appl Environ Microbiol. 2005 Dec;71(12):8228–35. doi: 10.1128/AEM.71.12.8228-8235.2005. PMID: 16332807

63. Vázquez-Baeza Y, Pirrung M, Gonzalez A, Knight R. EMPeror: a tool for visualizing high-throughput microbial community data. GigaScience. 2013 Nov 26;2(1):16. doi: 10.1186/2047-217X-2-16. PMID: 24280061

